# Lagomorph cranial biomechanics and the functional significance of the unique fenestrated rostrum of leporids

**DOI:** 10.1101/2024.11.21.624709

**Authors:** Amber P Wood-Bailey, Alana C Sharp

## Abstract

The crania of leporid lagomorphs are uniquely fenestrated, including the posterior cranial bones and the lateral portion of the maxilla. The functional significance of the highly fenestrated rostrum has received considerably little attention, despite being absent in other mammalian herbivores with a long rostrum. This unique feature is of particular interest when considering functional relationships between the loading regime and cranial structure. Two primary hypotheses have been suggested: maxillary fenestrations may be associated with the transmission and redirection of incisal occlusal forces, or fenestrations may reduce skull weight to assist with manoeuvrability and increase running speed. Here we apply a comparative approach using finite element analysis to determine how the overall stress and strain environment is affected by the presence or absence of maxillary fenestrations. We compare three lagomorph species with various degrees of latticing in the fenestrated rostrum, with two macropods that do not have fenestrations. We then produce theoretical models of the three lagomorphs by filling in the fenestrated region. Our results show that the presence of fenestrations makes little difference to the overall stress experienced through the cranium and does not impact the efficiency of incisor biting. This adds to the increasing evidence that features of lagomorph cranial morphology correlate with locomotor demands, adapting to loads other than mastication. Modulating cranial mass with fenestrations may provide benefits of a lighter skull, while still providing enough surface area for muscle attachments.

## Introduction

The leporid lagomorphs (rabbits and hares) display distinctive cranial morphology that distinguish them from other mammalian taxa. Among these are pronounced ventral facial tilting, potential cranial kinesis at an intracranial joint, and extensive fenestrations, particularly in the lateral wall of the maxilla. Such adaptations are thought to offer functional advantages associated with their characteristic rapid locomotion (DuBrul, 1950; Moss and Feliciano, 1977; Bramble, 1989; Kraatz et al., 2015; Watson et al., 2021). These unique features are of particular interest when considering functional relationships between the loading regime and cranial structure. Although cranial bone mass is typically regulated in response to masticatory loads, some regions may adapt to infrequent traumatic loads not related to feeding (Farke, 2008), or other functionally significant loads such as locomotion. Extensive fenestrations of mammalian cranial bones are uncommon, which has led to numerous hypotheses for their function in Leporidae.

Moss and Feliciano (1977) hypothesised that rostral fenestrations are associated with the transmission of incisal occlusal forces away from the lateral part of the maxilla, and instead posteriorly along the superior and inferior struts, which may be linked to the prominent downward facial tilt. They suggest that with an increasing angle of the ramus of the mandible in some leporids, incisal forces are redirected along the dorsal and ventral struts, and away from the lateral plate. However, DuBrul (1950) originally suggested that the latticing of the maxilla could increase the efficiency of speedy locomotion by reducing weight of the rostrum. Recent biomechanical analyses suggest that these fenestrations may indeed optimise cranial weight reduction without compromising structural integrity (Watson et al., 2021), aligning with DuBrul’s hypothesis.

Extensive fenestrations in the lateral maxilla also vary morphologically between species and individuals. In the non-leporid lagomorphs, the pikas, the trait is expressed as one large vacuity. In some genera (such as *Nesolagus, Caprolagus* and *Oryctolagus*), the trait is found only above the bony remnant of the lacrimal duct (Wood-Bailey et al., 2022). In its most extreme form, fenestrations are found both above and below the bony remnant of the lacrimal duct in groups such as *Lepus, Sylvilagus* and *Pronolagus* (Wood-Bailey et al., 2022). Furthermore, this appears to reduce the weight of the skull relative to head mass in the species with extensive fenestrations, with the genus *Lepus* showing less isometric scaling between skull length and weight than the genus *Oryctolagus* (Bramble, 1989). The variation of maxillary fenestration appears correlated with locomotor capabilities across taxa, with faster-moving species generally exhibiting more extensive fenestrations (Bramble, 1989; Wood-Bailey et al., 2022).

The extent to which masticatory loads or locomotor loads influence cranial fenestrations is difficult to test experimentally through strain gauge measurements, but can be investigated through finite element (FE) modelling, an *in silico* technique that can predict how a structure will deform under certain loading conditions (Rayfield, 2007). FE modelling has been used to analyse the cranial stresses and strains of differing taxa during mastication (e.g., Sharp, 2015; Tseng and Flynn, 2015; DeSantis et al., 2020; Dutel et al., 2021; Smith et al., 2021; Watson et al., 2021; Cox and Watson, 2024), as cranial form is largely influenced by cyclic masticatory loads. Comparisons between leporids and rodents are common, based on both having large diastema and hypselodont dentition (DuBrul, 1950; Kraatz et al., 2021). However, here we compare with macropods (wallabies and kangaroos), also having a superficially similar cranial shape with a long rostrum and large diastema, yet lack the unique fenestrations of leporids. Using FE modelling we can manipulate the geometry of digital models to test functional implications of the presence or absence of maxillary fenestrations to explore the relationship between cranial morphology and biomechanical function, providing further insight into the adaptive significance of these unique cranial adaptations. By comparing leporid and morphologically similar non-leporid taxa, we aim to expand knowledge on the form-function relationship and how variations in cranial structure support ecological adaptations and high-speed locomotion in leporid lagomorphs.

## Materials and Methods

### Specimens

An adult European wild rabbit (*Oryctolagus cuniculus*) skull was scanned using X-Tek HMX 160 micro-CT scanner (X-Tek Systems Ltd, United Kingdom), at a resolution of 28 µm in each direction. A skull of an adult hispid hare (*Caprolagus hispidus*) from the Liverpool World Museum (NML 15.5.60.29) was microCT scanned at the University of Liverpool Centre for Preclinical Imaging using a PerkinElmer Quantum GX micro-CT scanner (120 μm voxel size, 90 kV, 88 μA). Micro-CT scans of a pika skull (*Ochotona princeps*) were downloaded from Morphosource (AMNH 120698, 23 µm voxels) (https://www.morphosource.org/). The skulls of a swamp wallaby (*Walabia bicolor*; NMV C10226, 199 μm voxel size) and a red kangaroo (*Macropus rufus*; NMV C23045; 209 μm voxel size) were CT scanned using a Siemens Sensation 64 scanner (Siemens Medical Solutions) at St. Vincent’s Public Hospital in Melbourne, Australia.

### Model construction and loading conditions

The scan data were loaded into the image visualisation software program Avizo (TermoFisher Scientific) for automated and manual segmentation, to generate three-dimensional (3D) surface models. The surface models were smoothed and edited to improve the quality of the mesh, including testing the aspect ratio and the dihedral angles of the surface triangles. The aspect ratios of the triangles were adjusted to below 10, the dihedral angles were set at above 10 degrees and the tetra-quality was below 25 to ensure a good quality mesh. The surface meshes were then converted to solid 3D FE meshes composed of 4-node tetrahedral elements (tet4) and exported as an Abaqus input file (*.inp) for easy importation to the FEA software package Abaqus 2022 (Simulia). Models consisted of 1.2 million to 5 million elements using consistent element sizes by controlling for edge length and adjusting for the model size.

The material properties of the bone were made homogenous and isotropic with values of Young’s modulus (17.5 GPa) and Poisson’s ratio (0.3) from recent material properties data in the rabbit skull (Wang et al., 2024). These values were assigned in order to enable a direct comparison among models.

Boundary conditions were assigned to the model to prevent rigid body motion. To not over constrain the model, the left temporomandibular joint (TMJ) was constrained against all displacement, and the right TMJ was constrained in the vertical and in the anterior–posterior directions only. Single nodes were placed at the centre of each incisor to represent a biting point and only constrained in the direction of occlusion.

The jaw muscles were modelled by applying the muscle forces over the entire surface of the origin of the masseter, temporalis, and medial pterygoid, with a local co-ordinate system to direct the force to their corresponding insertion on the mandible. To estimate the muscle force for *Oryctolagus*, physiological cross-sectional area (PCSA) was calculated for each muscle following dissection of a fresh wild European rabbit, and chemical digestion of the masticatory muscles for mass and fibre length (muscles dissected: dorsal superficial masseter, ventral superficial masseter, anterior deep masseter, posterior deep masseter, temporalis, and medial pterygoid). A combined muscle force for the masseter was calculated from these individual muscle forces. Our PCSA (and subsequent force) estimation for the temporalis is much lower than previous measurements (Watson et al., 2014; Watson et al., 2021), possibly because we were not able to accurately dissect the deep temporalis. Compared to closely related mammals (such as rodents), the leporid temporalis muscle is greatly reduced in size (Guelinckx et al., 1986; Bramble, 1989), so we believe our estimates reflect this significant reduction. To estimate the muscle forces for *Caprolagus*, the muscle input forces for *Oryctolagus* were scaled to body mass (Dumont et al., 2009; Strait et al., 2010). Body mass estimation values were taken from regression statistics for body mass as a function of skull length for rodents (Reynolds, 2002).

Ochotonids appear to have a much larger attachment area for temporalis compared to leporids (Bramble, 1989). Therefore, rather than scaling the muscles forces from our *Oryctolagus* dissections for the *Ochotona* model, muscle forces were estimated by scaling forces obtained from Watson et al. (2021). These more accurately represent the relatively larger temporalis in *Ochotona*, while still maintaining relative contributions from the masseter and pterygoids of lagomorphs, rather than scaling from a rodent model.

For the macropods, muscle forces were obtained from Sharp (2015). As with the lagomorphs, the individual muscle forces were combined to produce a single force for the masseter, temporalis and pterygoid muscles. These where then applied over the entire origin for each muscle and the force was directed to the insertion on the mandible.

### Analysis of model performance

To assess the structural performance of each model, we report a combination of stress and strain, and mechanical advantage. Mechanical advantage was calculated by dividing the bite force (reaction force) by the total input adductor muscle force. Comparisons of the distribution of von Mises (VM) stress and maximum and minimum principal strain (primarily tensile and compressive strain respectively) are useful to infer ecological variation between species. To assess the overall spread of stress, the VM stress of each element was also exported from Abaqus for analysis in R. VM stress determines the potential for a material to yield. A higher value will result in an increased likelihood of a material fracturing. VM stress is useful because it combines tensile, compressive and shear stresses into a single value and is related to failure through ductile yielding as opposed to brittle fracture (Dumont et al., 2009), which is suitable for biological materials such as bone. However, reaching the failure point of a skull during maximal biting is unlikely in biological structures, so comparing the absolute values of VM stress are not as informative as identifying differences between the models.

## Results

The model of *Oryctolagus* was validated against in vivo bite forces (Watson et al., 2014) by comparing these with our computed bite reaction forces (69.1 ±13.3 N and 50.5 N respectively). Our other models could not be formally validated due to the lack of in vivo data and scarcity of specimens. However, for the present study, a comparative approach has been applied, which compares relative stress and strain values, and is not intended to predict absolute values. By applying the same modelling approach and material properties, we can assume, to some degree, that the other models are also within realistic values.

The mechanical advantage of biting provides a scale independent estimate of the efficiency at which the muscle force is translated to bite force, which is driven by the jaw lever system and the length from the fulcrum (TMJ) to the bite point. For incisor biting, *Macropus* had the lowest mechanical advantage (0.14) and *Caprolagus* had the highest (0.27). There was very little change after filling the rostral fenestrations: the mechanical efficiency of *Oryctolagus* went down from 0.20 to 0.19; *Caprolagus* went up from 0.27 to 0.32; and *Ochotona* remained the same at 0.20 for both.

The distribution of stress and strain across the models show some clear differences, but with some common similarities. The *Ochotona* and *Oryctolagus* models had the highest VM stress (Figure 1). This was concentrated along the zygomatic arch (particularly at the TMJ), rostrum, and around the orbit. Along the zygomatic arch, high stress is observed due to compression of the superior border and tension on the inferior border of the zygomatic arch (Figure 2 and 3). Lower stress is observed along the zygomatic arch in *Caprolagus, Macropus* and *Wallabia*. In *Ochotona*, high stress is also recorded around the orbit (20-30 MPa). This is predominantly due to tensile strains at the anterior border and compressive strain superior and posterior of the orbit. High stress is also observed on the anterior rostrum of *Oryctolagus, Macropus* and *Wallabia*, just superior to the incisors. This is due to high compressive strains (Figure 3).

**Figure 1.**
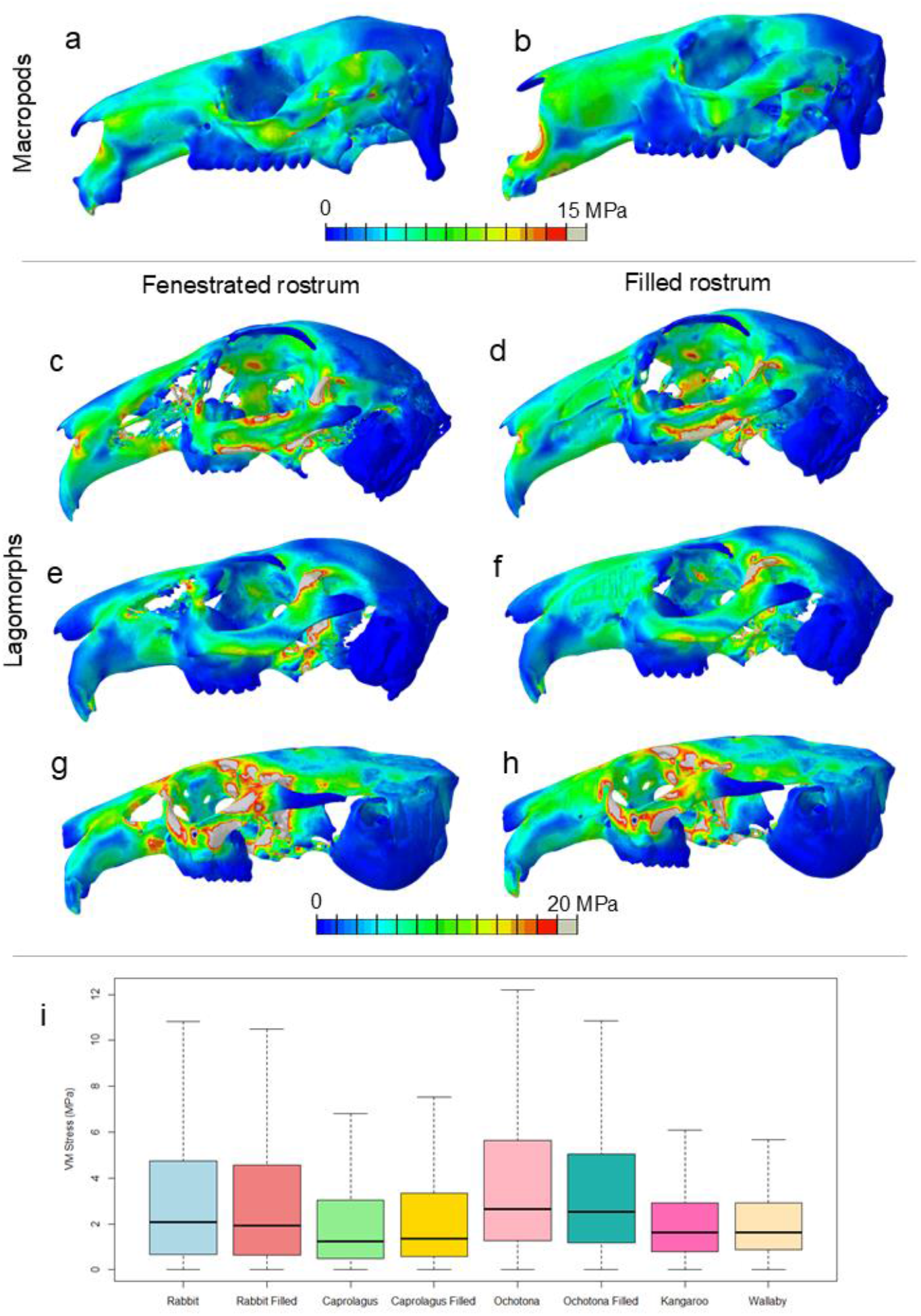
Von Mises stress distribution maps for macropods (a-b) and Lagomorphs (c-h). Low values are coloured in blue and high values are red; grey is beyond the maximum set with the scale. Due to lower values in the macropods, two different scales had to be used for a more accurate visualization of the data. Boxplots (i), scaled to skull surface area, show the spread of stress over the entire model with outliers removed. (a) *Walabia bicolor*, (b) *Macropus rufus*, (c-d) *Oryctolagus cuniculus*, (e-f) *Caprolagus hispidus*, (g-h) *Ochotona princeps*.

**Figure 2.**
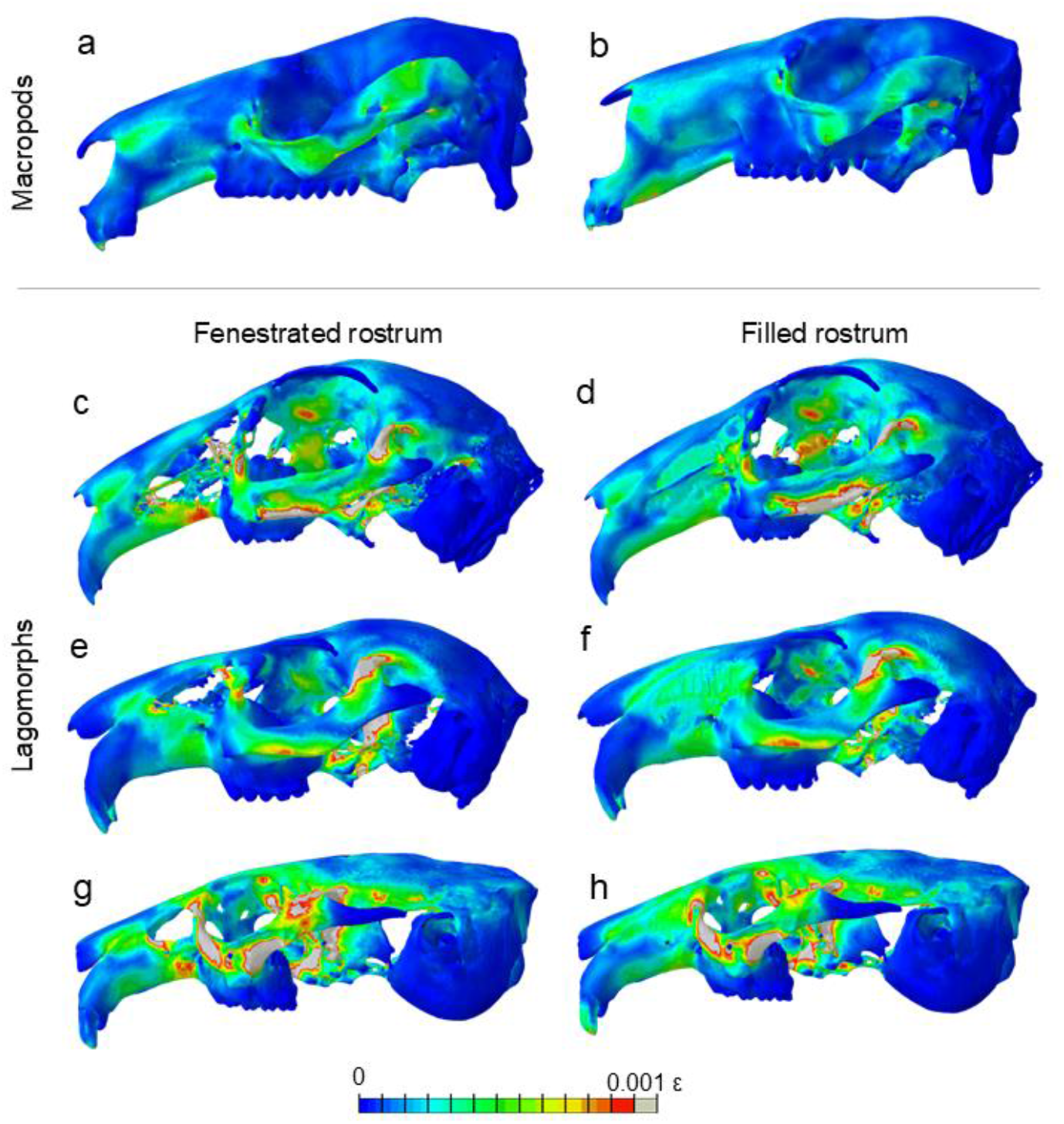
Minimum principal strain (predominantly tension) distribution maps for macropods (a-b) and Lagomorphs (c-h). Low values of tension are coloured in blue and high values are red; grey is beyond the maximum set with the scale. (a) *Walabia bicolor*, (b) *Macropus rufus*, (c-d) *Oryctolagus cuniculus*, (e-f) *Caprolagus hispidus*, (g-h) *Ochotona princeps*.

**Figure 3.**
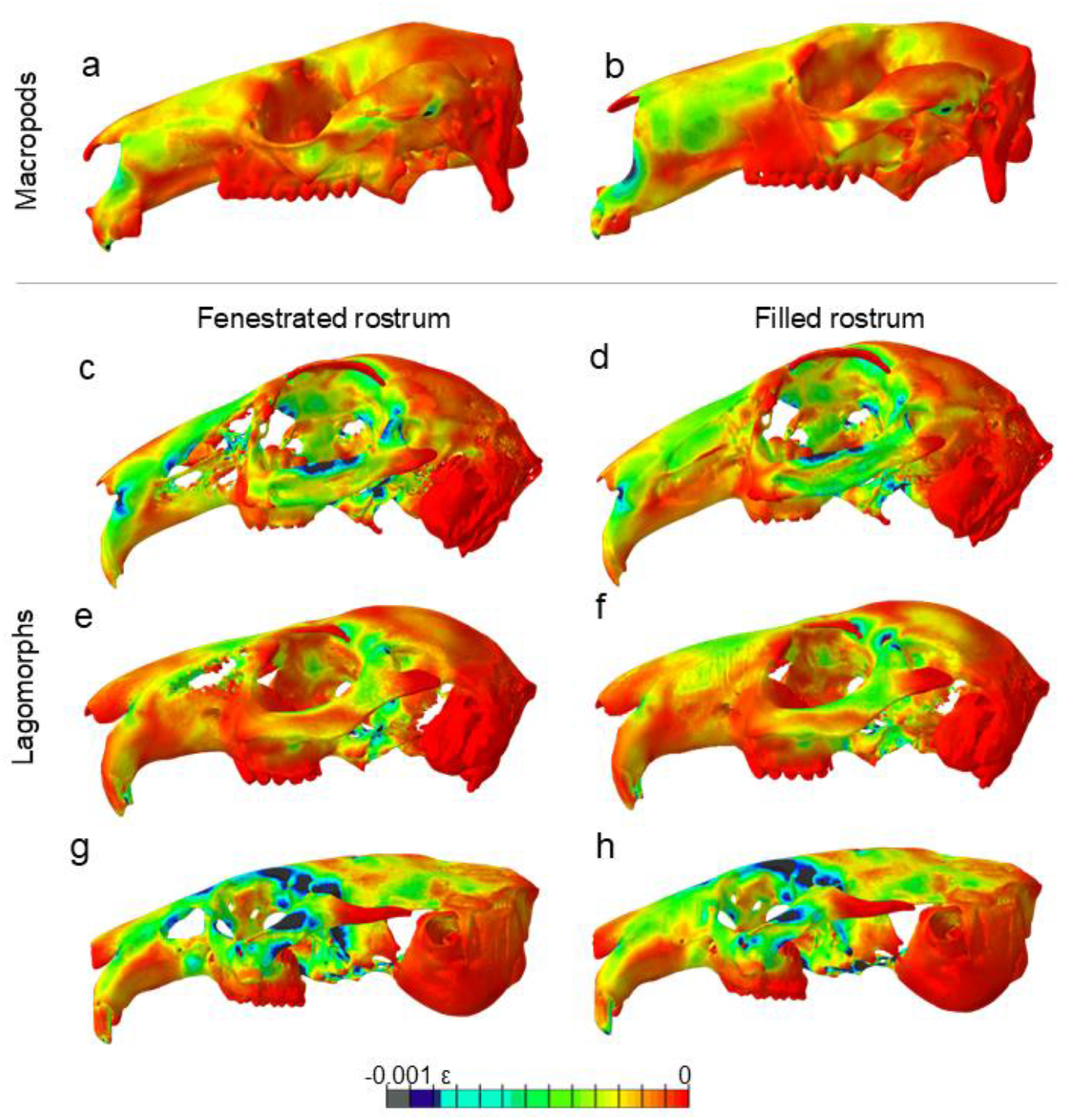
Maximum principal strain (predominantly compression) distribution maps for macropods (a-b) and Lagomorphs (c-h). Low values of compression are coloured in red and high values are blue-black; grey is beyond the maximum set with the scale. (a) *Walabia bicolor*, (b) *Macropus rufus*, (c-d) *Oryctolagus cuniculus*, (e-f) *Caprolagus hispidus*, (g-h) *Ochotona princeps*.

When comparing the fenestrated and filled models of *Oryctolagus, Caprolagus* and *Ochotona*, there is almost no difference in stress distribution posterior to the anterior border of the orbit (Figure 1). When filled, the lateral bone of the rostrum experiences stresses of similar magnitude to that in the kangaroo and wallaby (3-9 MPa), indicating that loads from incisor biting do propagate through the rostrum regardless of the presence of absence of fenestrations. The filled rostrums have lower overall stress compared to the fenestrated rostrums due to the occurrence of higher stress along the thin struts in *Oryctolagus*, and around the single large fenestration in *Ochotona* and *Caprolagus*.

## Discussion

This study compares leporid and morphologically similar non-leporid taxa to explore functional implications of the presence or absence of maxillary fenestrations. This unique feature, found in leporid lagomorphs but absent in other mammalian herbivores with a long rostrum, is particularly interesting when considering functional relationships between cranial loading and cranial structure. Our findings reveal that the fenestrated rostrum may be optimized for both cranial weight reduction and biomechanical performance during high-speed locomotion and feeding. In examining models of *Oryctolagus, Caprolagus*, and *Ochotona* with and without fenestrations, our results show that stress and strain distributions are consistent across filled and fenestrated conditions, especially posterior to the orbit. This suggests that while the fenestrations minimally influence the mechanical performance under biting loads, they may confer significant benefits for reducing skull weight, as proposed by DuBrul (1950) and further supported by recent studies (Watson et al., 2021; Wood-Bailey et al., 2022).

The similar stress distribution in fenestrated and filled models suggests that the fenestrations do not compromise structural integrity during biting, and may not function to redirect incisal loads along the dorsal and ventral struts of the rostrum as hypothesised by Moss and Feliciano (1977). Our comparison between lagomorphs and non-lagomorph taxa without fenestration, the macropods, indicates that the presence of fenestrations does not drastically alter the efficiency of bite force transmission or the distribution of cranial stress. Filling the fenestrations in *Oryctolagus, Caprolagus*, and *Ochotona* does reduce stress in the rostrum, which more closely resemble macropods without fenestrations; overall, models without fenestrations show a more uniform stress distribution across the rostrum. This pattern suggests that, while fenestrations do not offer an advantage during biting, it does not negatively impact leporids either.

Lagomorph species exhibiting more extensive fenestrations, such as *Lepus* and *Sylvilagus*, are typically associated with faster locomotion, while taxa with fewer or smaller fenestrations, such as *Caprolagus*, display relatively lower mechanical demands (Wood-Bailey et al., 2022). This distribution pattern suggests that fenestration degree correlates with locomotor demands, reinforcing the idea that leporid cranial morphology is adapting to loads other than mastication. The reduced mass in fenestrated species may facilitate rapid accelerations and agile manoeuvres necessary for predator avoidance, a critical selective pressure in open habitats with aerial predators where speed is essential for survival. Modulating cranial mass in this way may provide benefits of a lighter skull, while still providing enough surface area for muscle attachments.

The body size differences and distinct locomotor strategies between macropods and leporids may also provide insight into the unique presence of rostral fenestrations in leporids. Macropods, such as kangaroos and wallabies, utilize a bipedal bounding or hopping gait that allows for high speeds, but relies more on elastic energy storage within the tendons and limb musculature to propel their larger bodies forward efficiently (Biewener and Baudinette, 1995; Kram and Dawson, 1998; Roberts and Azizi, 2011). This bounding movement is energy-conservative, which may reduce the need for extensive cranial modifications, such as weight reduction. Conversely, leporids achieve rapid locomotion through a more flexible, quadrupedal bounding gait characterized by high acceleration and directional agility; traits that are likely essential in evading aerial predators in open habitats. Minimizing cranial mass could facilitate faster head movements and visual adjustments during these high-speed manoeuvres. The fenestrated skull may therefore represent an adaptation that allows leporids to maximize speed and manoeuvrability for their relatively small body size, enhancing their ecological fitness in environments where rapid, agile movements are critical for survival. This evolutionary divergence in cranial morphology likely reflects differing mechanical demands imposed by the distinct locomotor strategies of leporids and macropods.

More data on species diets and running speeds could help to provide much needed ecological data to elucidate evolutionary adaptations to these functional pressures. While FEA provides a robust method for estimating stress and strain in complex cranial structures, future studies could also benefit from *in vivo* or *ex vivo* data across a wider range of lagomorph species during feeding and running. Additionally, expanding FEA to simulate various loading conditions that lagomorphs might encounter in natural settings, such as impacts during evasive manoeuvres, could further elucidate the functional significance of fenestrations and other cranial adaptations in leporids. Studies that examine ontogenetic changes in fenestration and how they correspond to shifts in locomotor or dietary demands could also provide insight into the evolutionary trajectory of these adaptations.

In conclusion, the fenestrated rostrum in lagomorphs appears to be a specialized adaptation primarily for weight reduction rather than load redistribution during biting, contrasting with other herbivores with elongated rostra. This adaptation likely confers an evolutionary advantage in high-speed, cursorial locomotion, especially in open environments where agility and speed are essential. Our findings add to the growing body of evidence that leporid cranial morphology reflects a unique evolutionary response to the functional demands of both feeding and predator avoidance, exemplifying the dynamic interplay between form and ecological function in the evolution of cranial biomechanics.

## Author contributions

A.W.B. and A.C.S. designed the study. A.C.S. and A.W.B. created the volumetric models. A.C.S. performed the FEA simulations. A.C.S. analysed data. A.W.B. and A.C.S. contributed to writing the manuscript.

## Acknowledgments

We would like to thank the staff that facilitated the scanning of specimens at the different facilities (Centre for Preclinical Imaging, a Liverpool Shared Research Facility (LIV-SRF); St. Vincent’s Public Hospital, Melbourne; Viper High Performance Computing facility of the University of Hull), as well as the AMNH Mammalogy Department for providing access to the *Ochotona princeps* scans on Morphosource.

## Notes

### Competing Interest Statement

The authors have declared no competing interest.

